# Immunoinformatics Prediction of Epitope Based Peptide Vaccine Against Listeria Monocytogenes Fructose Bisphosphate Aldolase Protein

**DOI:** 10.1101/649111

**Authors:** Mustafa Elhag, Mustafa Abubaker, Nagla Mohammad Ahmad, Esraa Musa Haroon, Ruaa Mohamed Alaagib, Sahar Obi Abd albagi, Mohammed A. Hassan

**Affiliations:** Faculty of Medicine, University of Seychelles-American Institute of Medicine, Seychelles; Faculty of Medical Laboratory Sciences, Sudan University of Science and Technology, Sudan; Department of Medical Microbiology, Faculty of Medical Laboratories Sciences, Al-Neelain University, Sudan; Faculty of Medical Pharmacology, Al-Ahfad University, Sudan; Department of Pharmacies, National Medical Supplies Fund, Sudan; Faculty of Medical Laboratory Sciences, Al-Neelain University, Sudan; Department of Bioinformatics, DETAGEN Genetics Diagnostic Center, Kayseri, Turkey

**Keywords:** Immunoinformatics, Listeria Monocytogenes, fructose-1, 6-bisphosphate aldolase, Peptide vaccine, Epitope

## Abstract

*Listeria Monocytogenes* represents an important food-borne pathogen worldwide that can cause life-threatening listeriosis disease especially in pregnant women, fetuses, elderly people, and immuno-compromised individuals with high mortality rates. Moreover, no vaccine against it exists. This study predicts an effective epitope-based vaccine against Fructose 1,6 Bisphosphate Aldolase (FBA) enzyme of Listeria Monocytogenes using immunoinformatics approaches. The sequences were retrieved from NCBI and several prediction tests were conducted to analyze possible epitopes for B-cell, T-cell MHC class I and II. 3D structure of the promising epitopes was obtained. Two epitopes showed high binding affinity for B-cells, while four epitopes showed high binding affinity for MHCI and MHCII. The results were promising to formulate a vaccine with more than 98% population coverage. We hope that these promising epitopes serves as a preventive measure for the disease in the future and recommend invivo and invitro studies.

## INTRODUCTION

Bacteria of the genus *Listeria* that are widely distributed in the environment comprise a group of gram-positive, facultative anaerobe, non-sporulating rods [1]. It consists of ten different species with *Listeria Monocytogenes* commonly found in humans [2]. *Listeria Monocytogenes* represents an important food-borne pathogen worldwide, that can cause life-threatening listeriosis disease in the susceptible groups including pregnant women, fetuses, elderly people, and immuno-compromised individuals, with a a considerable mortality rate (20–30%) [3]. The disease includes, but not limited to sepsis, meningitis, encephalitis, spontaneous abortion, or fever and self-limiting gastroenteritis in a healthy adult [2]. Food-borne listeriosis has a global economic and health burden due to the wide spread of *Listeria Monocytogenes* in food and food processing environments [4]. It capable of proliferating in different stressful environmental conditions, including high salinity, low temperature (refrigerator), and a wide range of pH values [4]. Although the incidence of listeriosis is relatively rare compared to other food-borne illnesses, listeriosis accounts for approximately 19% of deaths among all food-borne illness [5]. Studies have shown that *Listeria Monocytogenes* is the third leading cause of death from food-borne illness in the United States, with approximately 260 deaths annually. Mortality rates with confirmed *Listeria Monocytogenes* infections are around 15% but can be higher depending on patient status and comorbidities [6]. *Listeria Monocytogenes* has 13 different serotypes based on a variety of flagella and surface antigens. However, there are only three serotypes (1/2a, 1/2b, 4a) that can cause disease in humans [2]. Mendonça et al., (2016) mentioned that the surface exposed antigens of fructose-1,6-bisphosphate aldolase (FBA) class II in *Listeria* species is the antigen target of the previously described a hybridoma-derived antibody (mAb-3F8). They reported FBA as a novel immunogenic surface target useful for the detection of the genus *Listeria* [7]. 30-kDa protein is a fructose-1,6-bisphosphate aldolase (FBA), an enzyme of the glycolytic pathway that catalyzes the cleavage of its substrate fructose-1,6-bisphosphate (FBP) into glyceraldehyde 3-phosphate (G3P) and dihydroxyacetone phosphate (DHAP). There are two main classes of FBA: class 1 is known to form tetramers, and is present mainly in higher eukaryotes, such as animals, plants, and algae; while class II can form many different multimers, and is present mainly in bacteria [8]. Class II FBA has been studied as a potential target of new antibiotics [9, 10] and as vaccine antigen [11]. Besides, many studies have shown that FBA may play a role in pathogenesis by interacting with host’s plasminogen [12, 13] or promoting adhesion to host’s cells [14, 15]. Thus, FBA is considered a moonlighting protein (protein with two or more dissimilar functions) in many species [16] and may have significant role in both physiology and pathogenesis.

Understanding of epitope/antibody interaction is the key to construct potent vaccines and effective diagnostics [17]. Most of immunoinformatics researches are stressed on the design and study of algorithms for mapping potential B- and T-cell epitopes that speed up the time and lower the costs needed for laboratory analysis of pathogen products. Using such tools and information (reverse vaccinology) to analyse the sequence areas with potential binding sites can lead to the development of new vaccines [18].

This is the first study to predict an effective epitope-based vaccine against FBA enzymes of *Listeria Monocytogenes* using immunoinformatics approaches.

## MATERIALS AND METHODS

### Protein Sequence Retrieval

A total of 1750 *Listeria monocytogenes* FBA strains were retrieved from National Center for Biotechnology Information (NCBI) database on March 2019. The protein sequence had length of 284 with name fructose-1,6-bisphosphate aldolase.

### Multiple Sequence Alignment

Using BioEdit Sequence Alignment Editor Software version 7.2.5, the retrieved sequences of *Listeria Monocytogenes* FBA were subjected to Multiple Sequence Alignment (MSA) using ClustalW to obtain the conserved regions and amino acid composition of the protein.[19] [20]

### Sequenced-Based Method

The reference sequence (WP_096833455.1) of *Listeria Monocytogenes* FBA was submitted to different prediction tools at the Immune Epitope Database (IEDB) analysis resource (http://www.iedb.org/). Epitope Analysis Resources were used to predict B and T cell epitopes. Conserved epitopes would be considered as candidate epitopes for B and T cells. [21]

### B Cell Epitope Prediction

Candidate epitopes were analysed using several B cell prediction methods from IEDB (http://tools.iedb.org/bcell/), to identify the surface accessibility, antigenicity and hydrophilicity. The Bepipred Linear Epitope Prediction 2 was used to predict linear B cell epitope with default threshold value -.012 (http://tools.iedb.org/bcell/result/). The Emini Surface Accessibility Prediction tool was used to detect the surface accessibility with default threshold value 1.000 (http://tools.iedb.org/bcell/result/). The Kolaskar and Tongaonker Antigenicity method was used to identify the antigenicity sites of candidate epitope with default threshold value 1.032 (http://tools.iedb.org/bcell/result/). The Parker Hydrophilicity Prediction tool was used to identify the hydrophilic, accessible, or mobile regions with default threshold value 1.695. [22-26]

### T Cell Epitope Prediction MHC Class I Binding

Analysis of peptide binding to the MHC (Major Histocompatibility complex) class I molecule was assessed by the IEDB MHC I prediction tool (http://tools.iedb.org/mhci/) to predict cytotoxic T cell epitopes. The presentation of peptide complex to T lymphocyte undergo several steps. Before the prediction, all human allele lengths were set to 9 amino acid. Artificial Neural Network (ANN) 4.0 prediction method was used to predict the binding affinity. The half-maximal inhibitory concentration (IC50) value required for all conserved epitope that bind to at allele at score less than 100 were selected. [27-33]

### T Cell Epitope Prediction MHC Class II Binding

Prediction of T cell epitopes and interaction with MHC Class II was assessed by the IEDB MHC II prediction tool (http://tools.iedb.org/mhcii/). Human allele references set were used to determine the interaction potentials of T cell epitopes and MHC Class II allele (HLA DR, DP and DQ). NN-align method was used to predict the binding affinity. IC50 values at score less than 500 were selected.[34-37]

### Population Coverage

In IEDB, the population coverage link was selected to analyse the epitopes. This tool calculates the fraction of individuals predicted to respond to a given set of epitopes with known MHC restrictions (http://tools.iedb.org/population/iedbinput). The appropriate checkbox for calculation was checked based on MHC I, MHC II separately and combination of both. [38]

### 3D Structures

3D structure was obtained using raptorX (http://raptorx.uchicago.edu) i.e. a protein structure prediction server developed by Xu group, excelling at predicting 3D structures for protein sequences without close homologs in the Protein Data Bank (PDB). USCF chimera (version 1.8) was the program used for visualization and analysis of molecular structure (http://www.cgl.uscf.edu/chimera). [39, 40]

## RESULTS

### Multiple Sequence Alignment

The conserved regions and amino acid composition for the reference sequence of *Listeria Monocytogenes* FBA is illustrated in figure 1 and 2 respectively.

### B-cell epitope prediction

The reference sequence of *Listeria Monocytogenes* FBA was subjected to Bepipred linear epitope, Emini surface accessibility, Kolaskar & Tongaonkar antigenicity and Parker hydrophilicity Prediction methods to test for various immunogenicity parameters (Table 1 and Figures 3-8). 3D structure of the proposed B cell epitopes is shown (Figure 7 & 8).

**Table 1:**
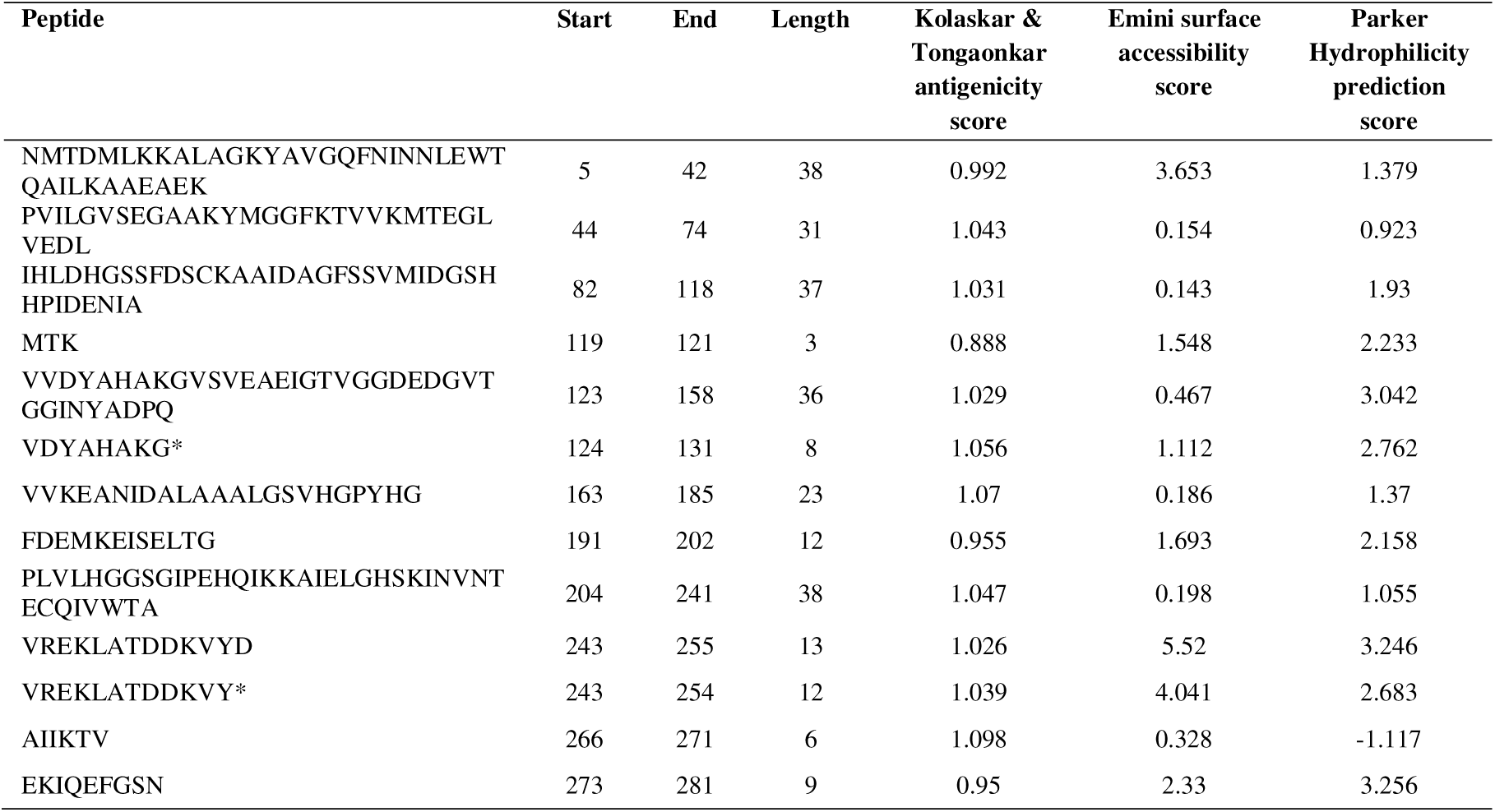
List of conserved peptides with their antigenicity, EMINI surface accessibility and Parker hydrophilicity scores. (* Peptides that successfully passed the three tests).

### Prediction of cytotoxic T-lymphocyte epitopes and interaction with MHC class I

The reference sequence was analyzed using (IEDB) MHC-1 binding prediction tool to predict T cell epitopes interacting with different types of MHC Class I alleles, based on Artificial Neural Network (ANN) with half-maximal inhibitory concentration (IC50) <100 nm. 61 peptides were predicted to interact with different MHC-1alleles. The most promising epitopes and their corresponding MHC-1 alleles are shown in (Table 2) followed by the 3D structure of the proposed T cell epitope (Figure 9).

**Table 2:**
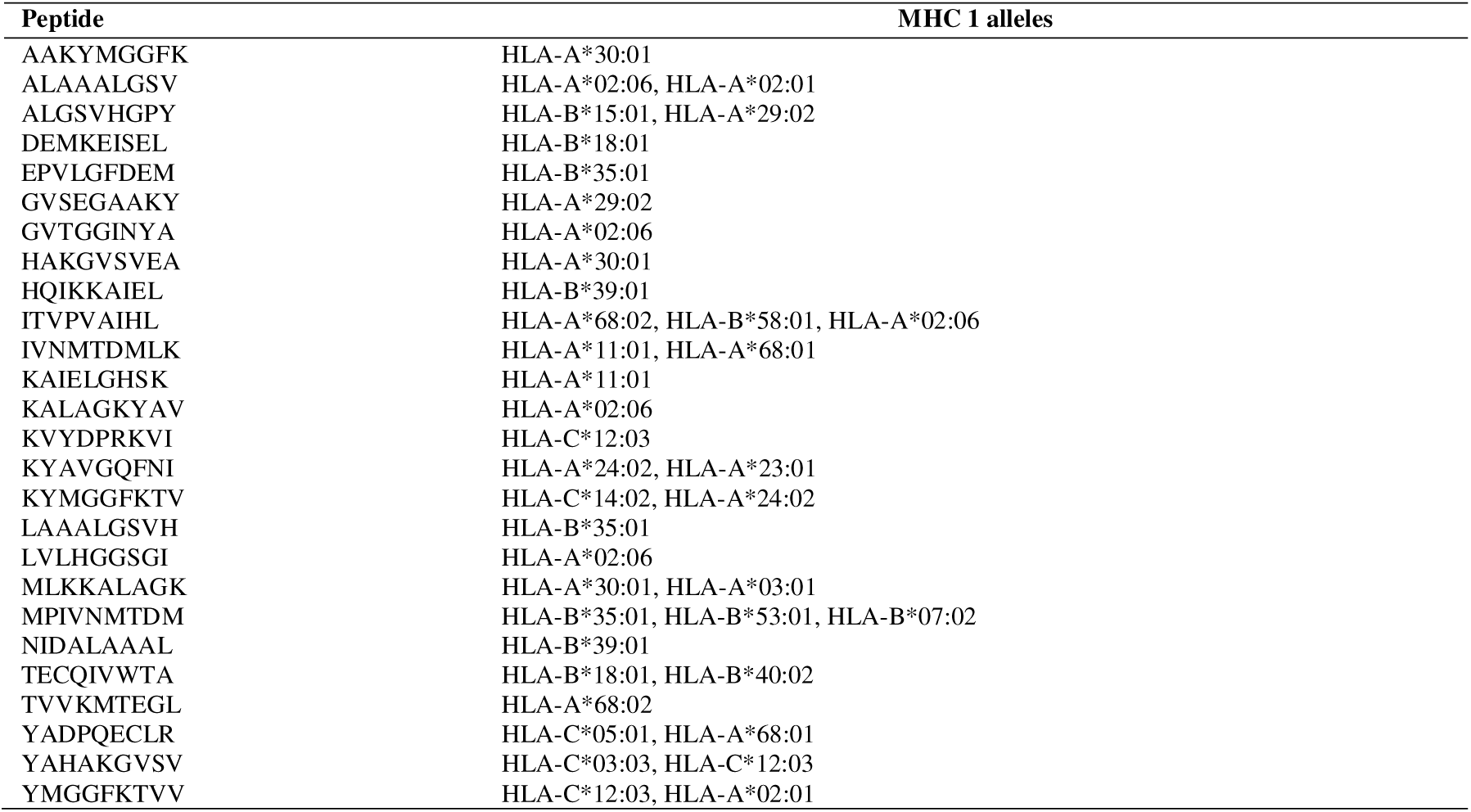
The most promising T cell epitopes and their corresponding MHC-1 alleles.

### Prediction of the T cell epitopes and interaction with MHC class II

Reference sequence was analyzed using (IEDB) MHC-II binding prediction tool based on NN-align with half-maximal inhibitory concentration (IC50) <500 nm; there were 388 predicted epitopes found to interact with MHC-II alleles. The most promising epitopes and their corresponding alleles are shown in (Table 3) along with the 3D structure of the proposed epitope (Figure 10).

**Table 3:** The most promising T cell epitopes and their corresponding MHC-2 alleles

| Peptide | MHC 2 alleles |
| --- | --- |
| VVDYAHAKGVSVEAE | HLA-DRB1*07:01, HLA-DQA1*05:01/DQB1*03:01, HLA-DRB1*01:01, HLA-DRB1*09:01, HLA-DRB5*01:01, HLA-DRB1*13:02, HLA-DQA1*03:01/DQB1*03:02, HLA-DRB1*1 |
| MTDMLKKALAGKYAV | HLA-DRB5*01:01, HLA-DRB1*01:01, HLA-DRB1*11:01, HLA-DRB1*09:01, HLA-DQA1*05:01/DQB1*03:01, HLA-DRB1*03:01, HLA-DRB1*15:01, HLA-DRB1*13:02, HLA-D |
| NMTDMLKKALAGKYA | HLA-DRB1*09:01, HLA-DRB5*01:01, HLA-DRB1*11:01, HLA-DRB1*01:01, HLA-DQA1*05:01/DQB1*03:01, HLA-DRB1*03:01, HLA-DRB1*15:01 |
| TDMLKKALAGKYAVG | HLA-DRB1*11:01, HLA-DRB5*01:01, HLA-DRB1*01:01, HLA-DQA1*05:01/DQB1*03:01, HLA-DRB1*09:01, HLA-DRB1*03:01, HLA-DRB1*15:01, HLA-DRB1*13:02, HLA-D |

### Population Coverage Analysis

All promising MHC I and MHC II epitopes were assessed for population coverage against the whole world.

**Table 4:** The population coverage of whole world for the most promising epitopes for MHC I, MHC II and MHC I& II combined.

| Country | MHC I | MHC II | MHC I&II combined |
| --- | --- | --- | --- |
| World | 94.15 % | 80.8 % * | 98.94 % * |

In population coverage analysis of MHC II; 12 alleles were not included in the calculation, therefore the above (*) percentages are for epitope sets excluding these alleles: HLA-DQA1*05:01/DQB1*03:01, HLA-DQA1*01:02/DQB1*06:02, HLA, HLA-DPA1*01:03/DPB1*02:01, HLA-DQA1*05:01/DQB1, HLA-DQA1*03:01/DQB1*03:02, HLA-DRB3*01:01, HLA-DRB4*01:01, HLA-DRB5*01:01, HLA-DQA1*05:01/DQB1*02:01, HLA-DPA1*03:01/DPB1*04:02, HLA-DQA1*04:01/DQB1*04:02, HLA-DPA1*02:01/DPB1*01:01.

## DISCUSSION

Different proposed peptides that can be recognized by B cell and T cell to produce antibodies against FBA of *Listeria Monocytogenes* were presented for the first time. Peptide vaccines overcome the side effects of conventional vaccines.

The reference sequence of *Listeria Monocytogenes* FBA was subjected to Bepipred linear epitope prediction 2 test, Emini surface accessibility test and Kolaskar and Tongaonkar antigenicity test and Parker hydrophilicity test in IEDB, to determine the binding to B cell and to test the immunogenicity and hydrophilicity respectively. Out of the five predicted epitopes using Bepipred 2 test, only two epitopes passed the other three tests (**VDYAHAKG, VREKLATDDKVY**).

26 epitopes were predicted to interact with MHCI alleles with IC50 < 100. Four of them were most promising (**ITVPVAIHL, MPIVNMTDM, MLKKALAGK, YMGGFKTVV**). 134 predicted epitopes were interacted with MHCII alleles with IC50 < 500. Four of them were most promising (**VVDYAHAKGVSVEAE, MTDMLKKALAGKYAV, NMTDMLKKALAGKYA, TDMLKKALAGKYAVG**). Eleven epitopes (**YAHAKGVSV, ALAAALGSV, TVVKMTEGL, MLKKALAGK, ITVPVAIHL, HAKGVSVEA, IVNMTDMLK, ALGSVHGPY, KALAGKYAV, NIDALAAAL, LAAALGSVH**) were shared between MHC I and II.

Excluding certain alleles for MHCII, population coverage for the most promising epitopes covered 98.94% of the whole world (MHCI & MHCII combined).

Many studies predicted peptide vaccines for different microorganisms such as, Rubella, Ebola, Dengue, Zika, HPV, Lagos rabies virus, and mycetoma using immunoinformatics tools. [41-50] Limitations include the exclusion of certain HLA alleles for the MHC II. We hope that the whole world will benefit from this epitope-based vaccine upon its successful development following invivo and invitro studies to prove it’s effectiveness.

## CONCLUSION

Peptide vaccines overcome the side effects of conventional vaccines. We presented different peptides that can produce antibodies against FBA of *Listeria Monocytogenes* for the first time. Two B cell epitopes passed the antigenicity, accessibility and hydrophilicity tests. Four MHCI epitopes were the most promising ones, while four for MHC II. Eleven epitopes were shared between MHC I and II. For the population coverage, the epitopes covered 98.94% of the alleles worldwide excluding certain ones.

---

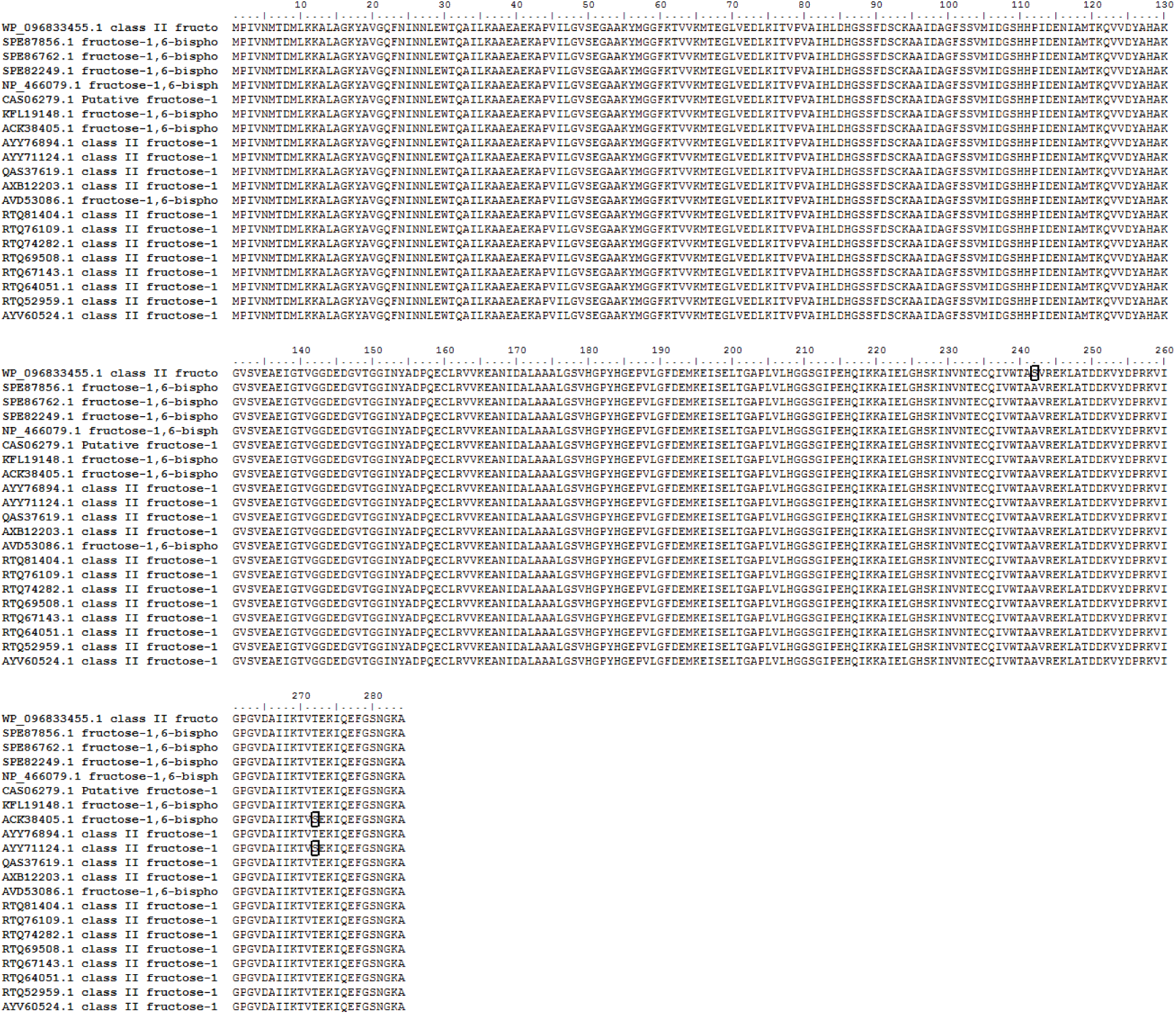

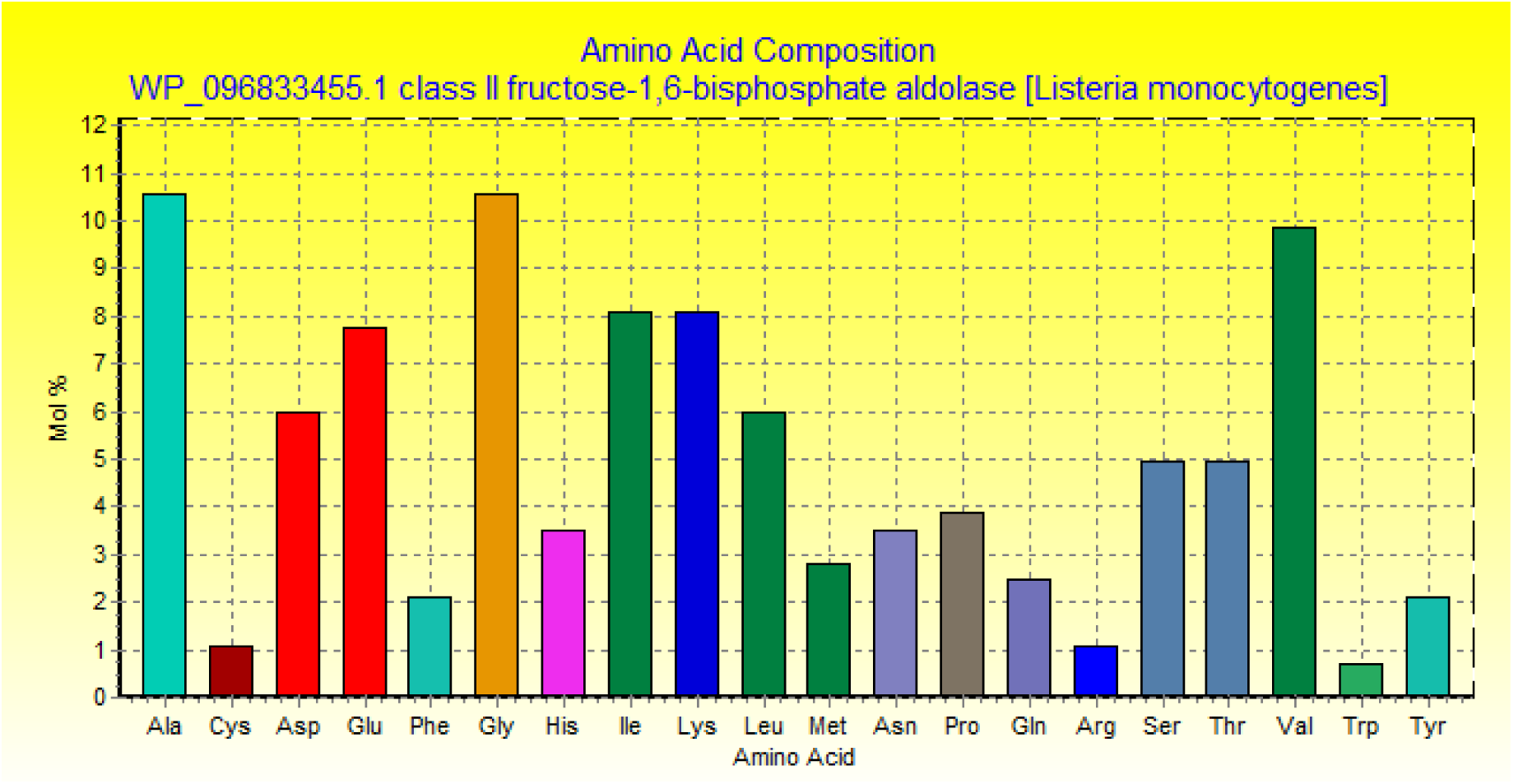

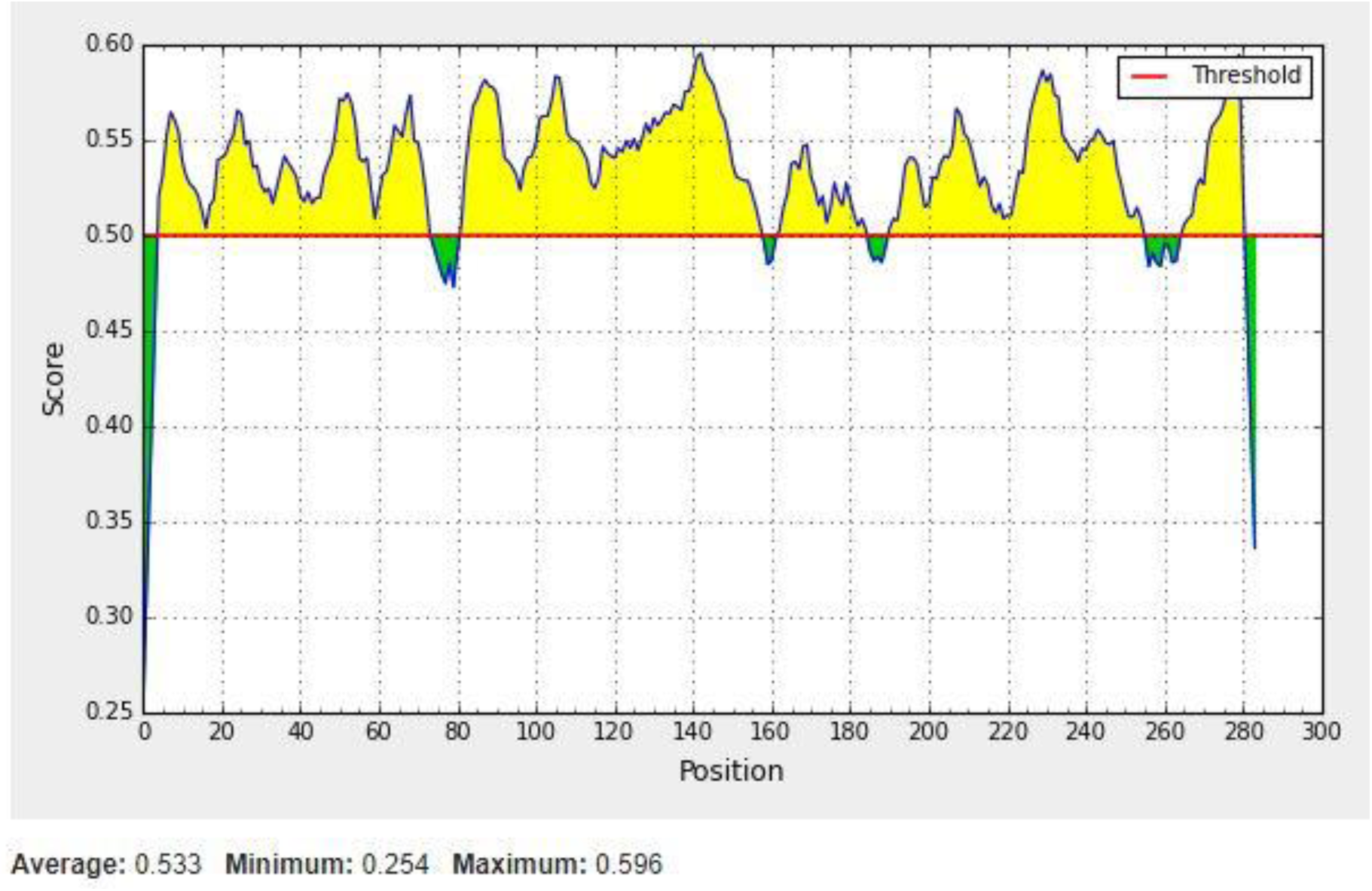

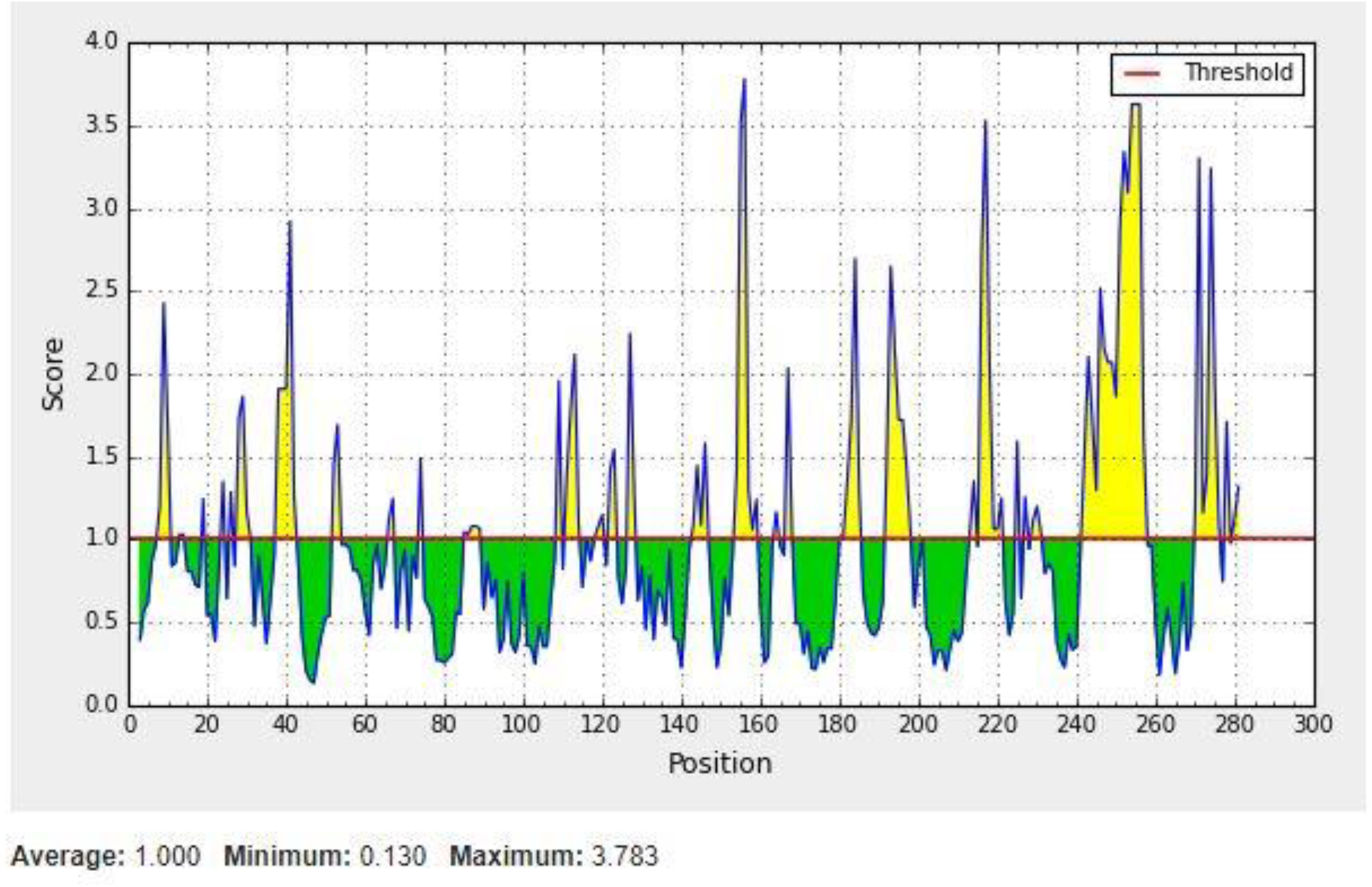

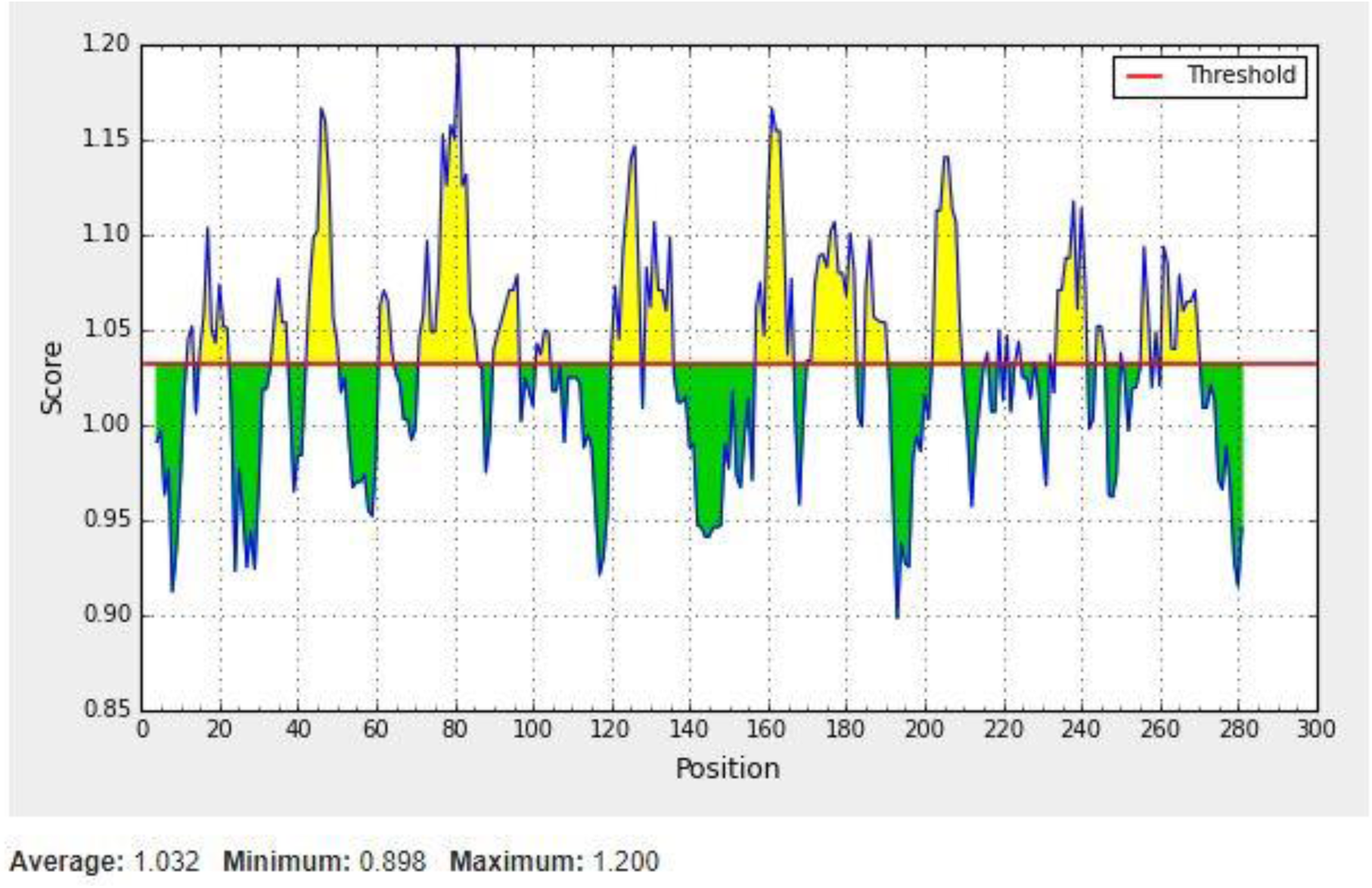

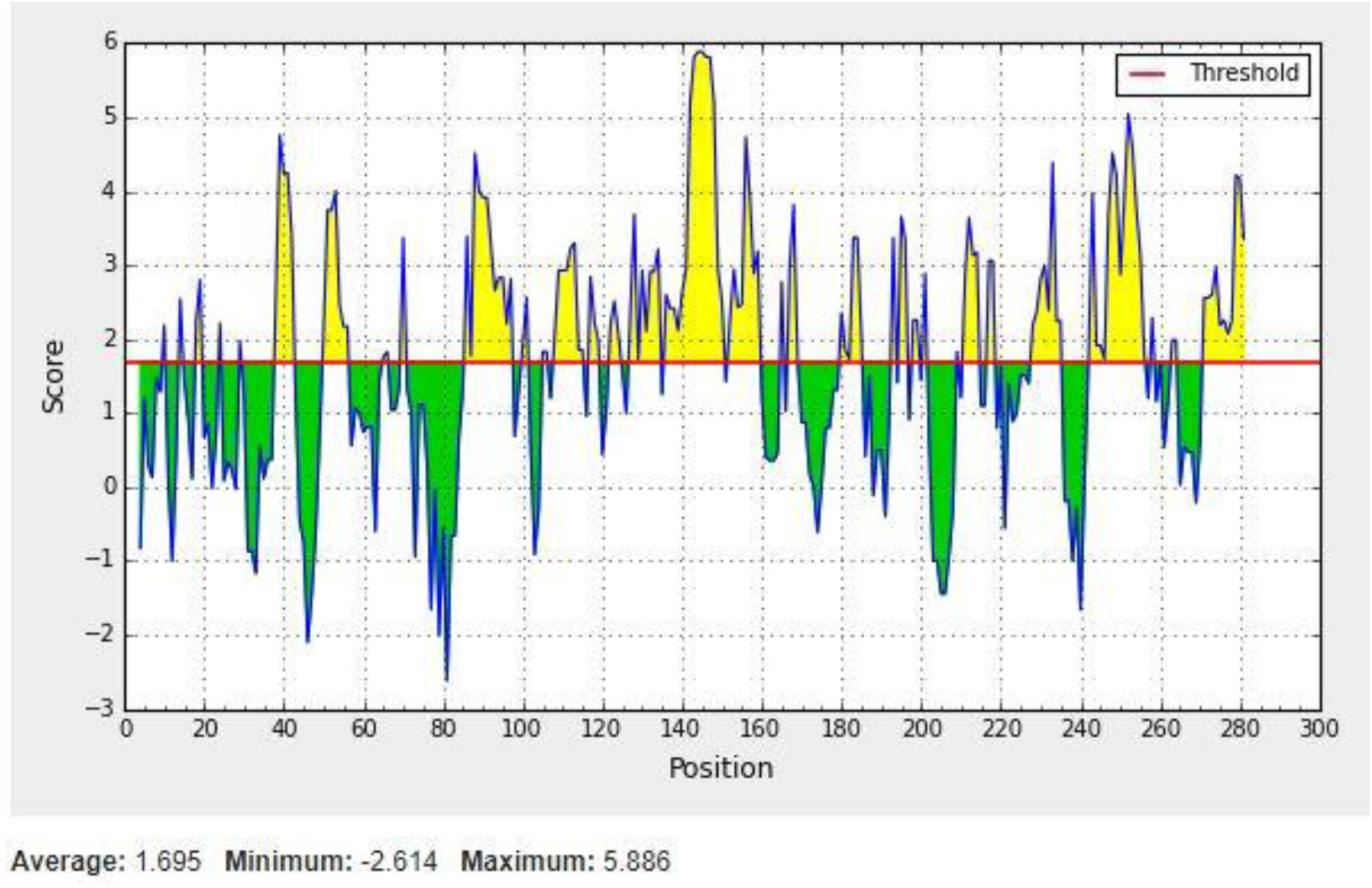

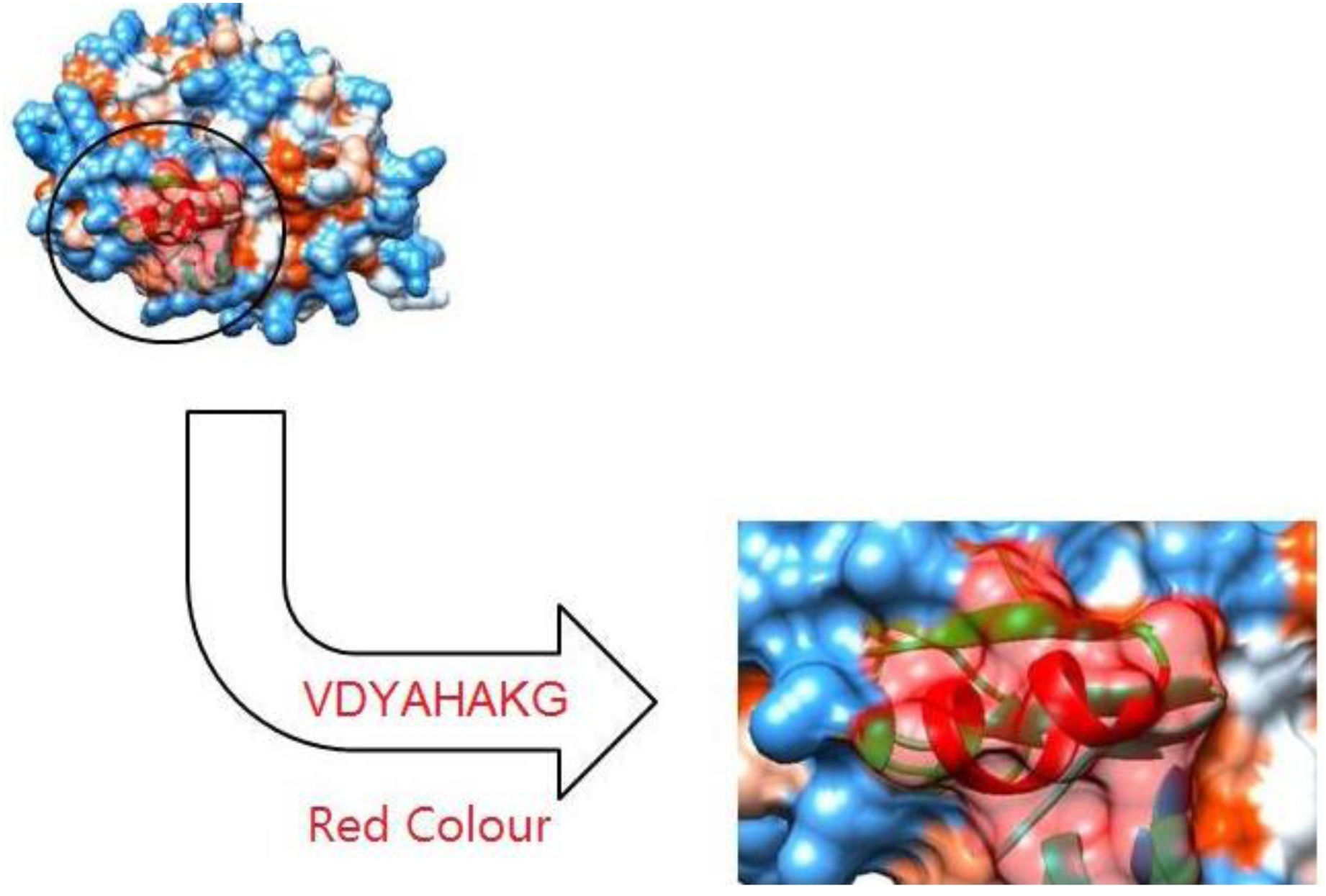

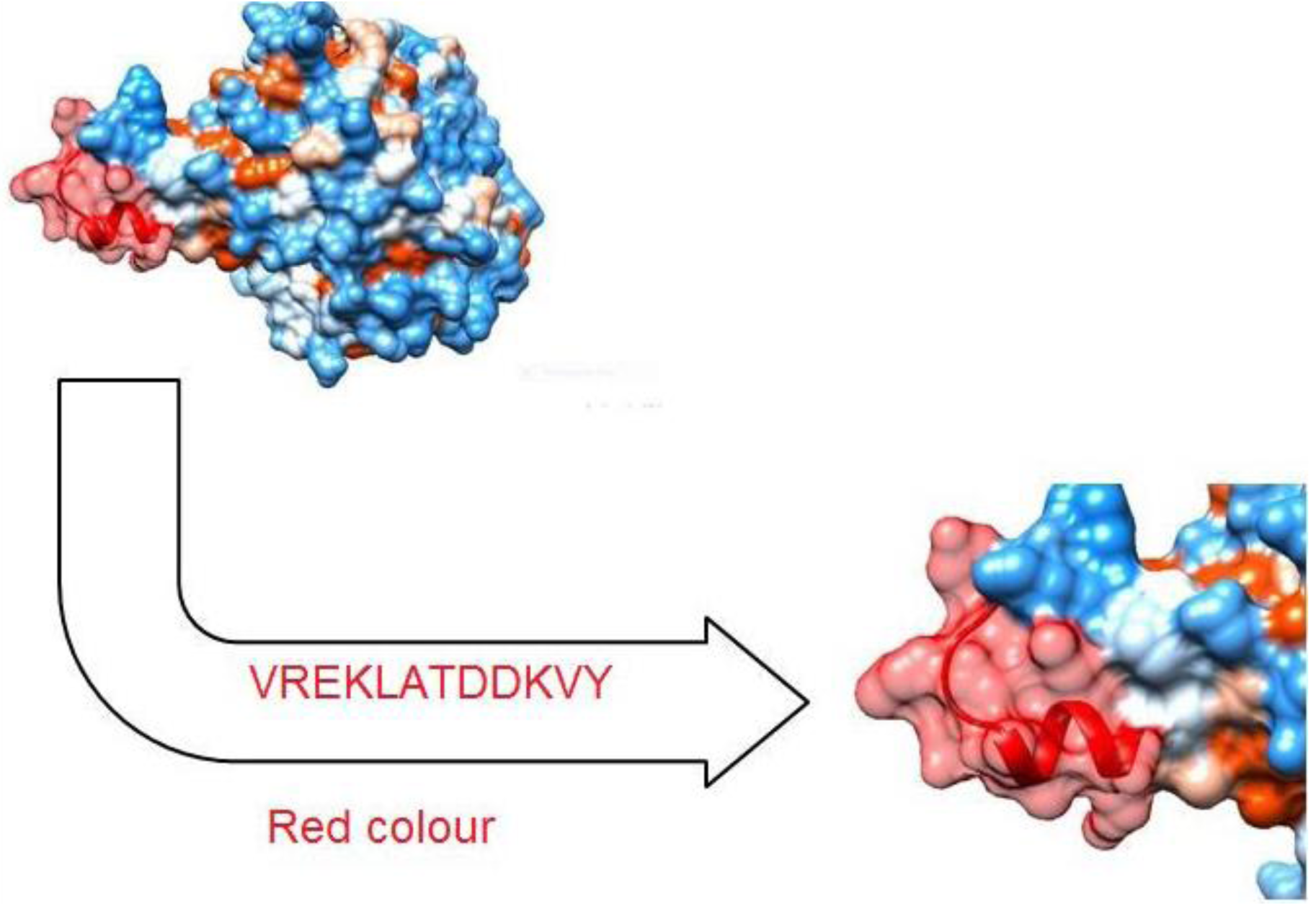

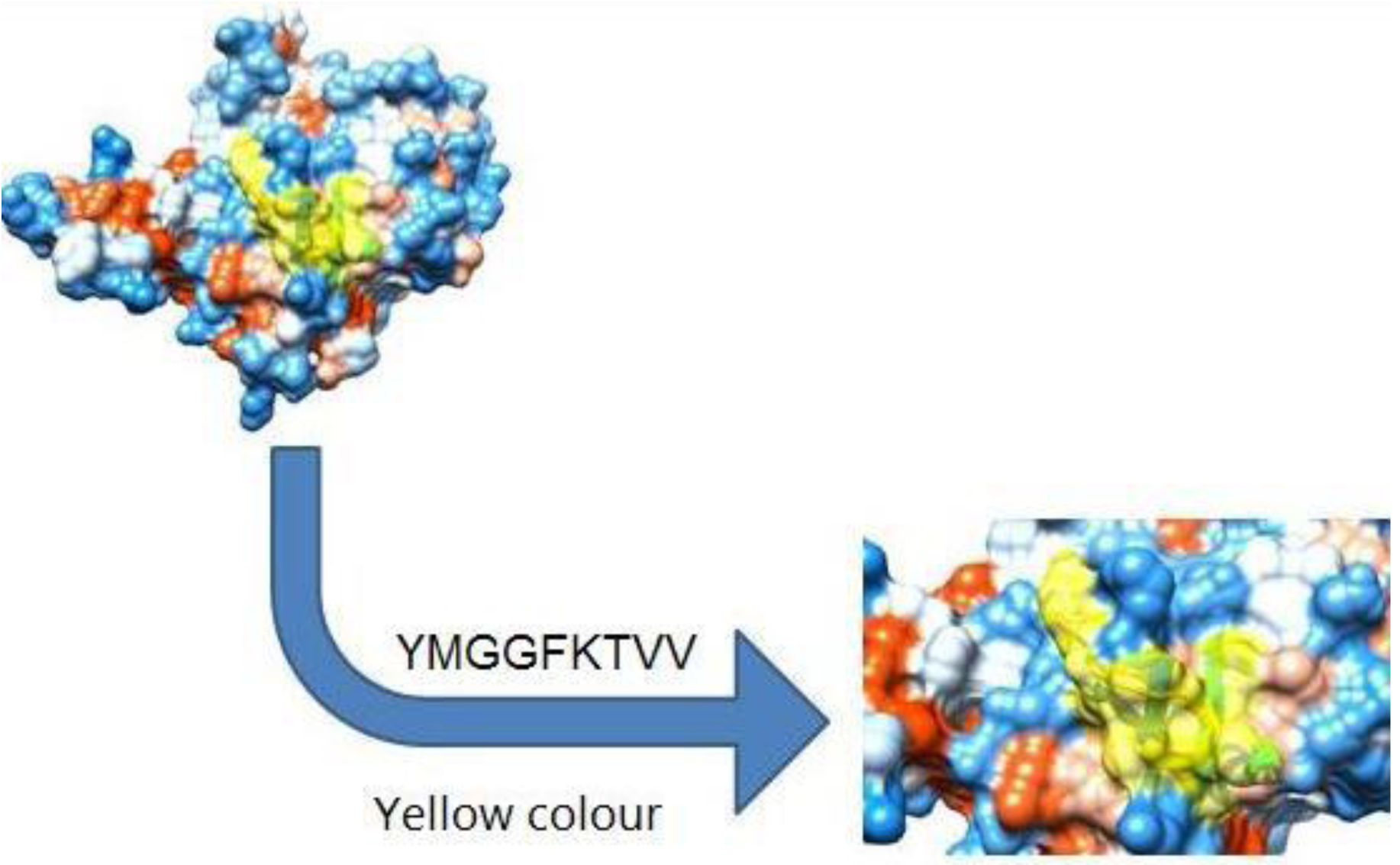

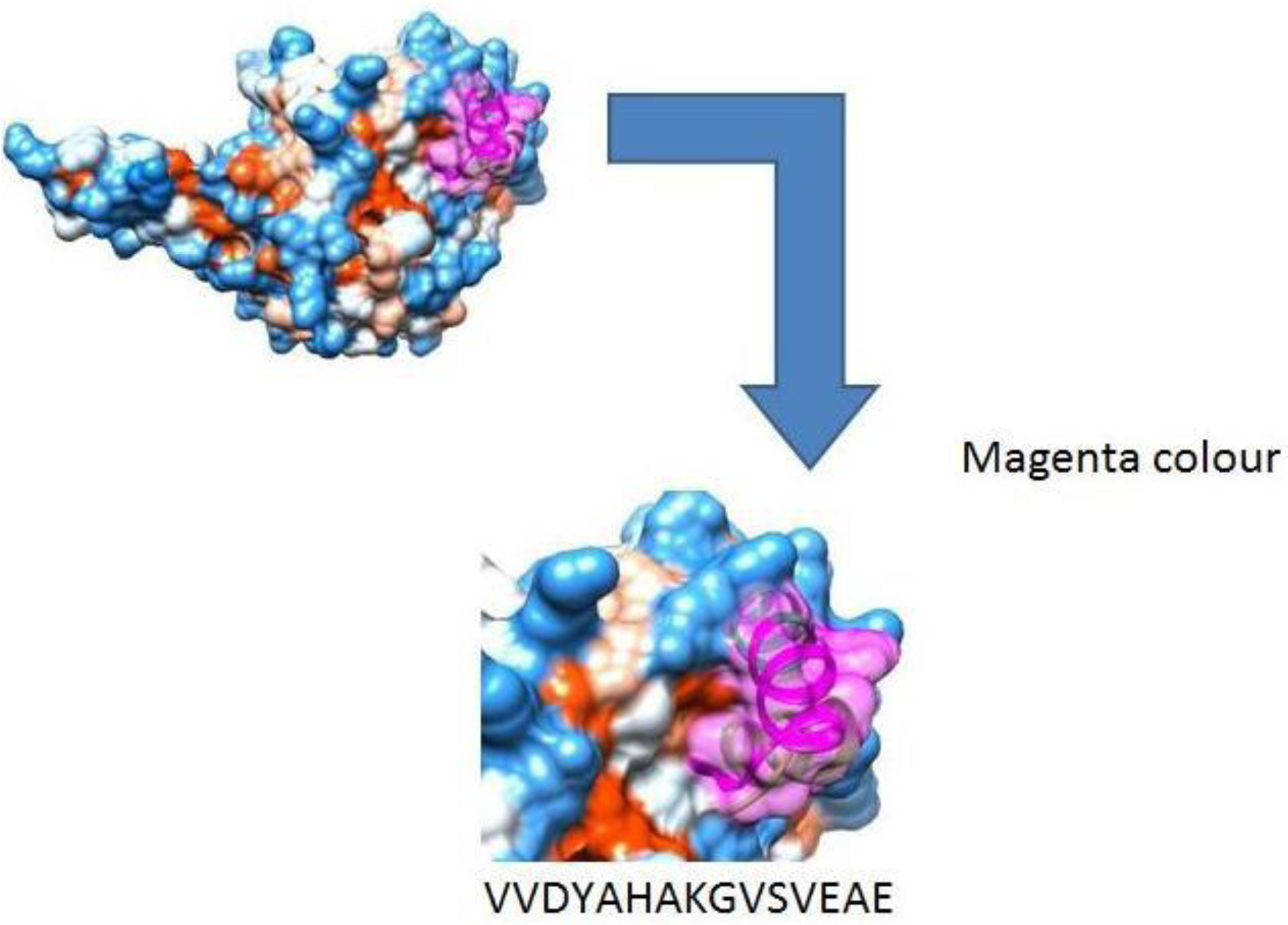

## References

1. Hain, T., et al., Pathogenomics of Listeria spp. International Journal of Medical Microbiology, 2007. 297(7-8): p. 541–557.

2. Hunt, K., et al., Challenge Studies to Determine the Ability of Foods to Support the Growth of Listeria monocytogenes. Pathogens, 2018. 7(4): p. 80.

3. Lomonaco, S., et al., Listeria monocytogenes in Gorgonzola: subtypes, diversity and persistence over time. International journal of food microbiology, 2009. 128(3): p. 516–520.

4. Chen, M., et al., Occurrence, Antibiotic Resistance, and Population Diversity of Listeria monocytogenes Isolated From Fresh Aquatic Products in China. Frontiers in microbiology, 2018. 9.

5. Scallan, E., et al., Foodborne illness acquired in the United States—major pathogens. Emerging infectious diseases, 2011. 17(1): p. 7.

6. Jackson, K.A., et al., Listeriosis outbreaks associated with soft cheeses, United States, 1998–2014. Emerging infectious diseases, 2018. 24(6): p. 1116.

7. Mendonça, M., et al., Fructose 1, 6-bisphosphate aldolase, a novel immunogenic surface protein on Listeria species. PloS one, 2016. 11(8): p. e0160544.

8. Katebi, A.R. and R.L. Jernigan, Aldolases utilize different oligomeric states to preserve their functional dynamics. Biochemistry, 2015. 54(22): p. 3543–3554.

9. Fonvielle, M., et al., Synthesis and biochemical evaluation of selective inhibitors of class II fructose bisphosphate aldolases: towards new synthetic antibiotics. Chemistry–A European Journal, 2008. 14(28): p. 8521–8529.

10. Daher, R., et al., Rational design, synthesis, and evaluation of new selective inhibitors of microbial class II (zinc dependent) fructose bis-phosphate aldolases. Journal of medicinal chemistry, 2010. 53(21): p. 7836–7842.

11. Elhaik Goldman, S., et al., Streptococcus pneumoniae fructose-1, 6-bisphosphate aldolase, a protein vaccine candidate, elicits Th1/Th2/Th17-type cytokine responses in mice. International journal of molecular medicine, 2016. 37(4): p. 1127–1138.

12. de la Paz Santangelo, M., et al., Glycolytic and non-glycolytic functions of Mycobacterium tuberculosis fructose-1, 6-bisphosphate aldolase, an essential enzyme produced by replicating and non-replicating bacilli. Journal of Biological Chemistry, 2011. 286(46): p. 40219–40231.

13. Chaves, E.G.A., et al., Analysis of Paracoccidioides secreted proteins reveals fructose 1, 6-bisphosphate aldolase as a plasminogen-binding protein. BMC microbiology, 2015. 15(1): p. 53.

14. Blau, K., et al., Flamingo cadherin: a putative host receptor for Streptococcus pneumoniae. The Journal of infectious diseases, 2007. 195(12): p. 1828–1837.

15. Tunio, S.A., et al., The moonlighting protein fructose-1, 6-bisphosphate aldolase of Neisseria meningitidis: surface localization and role in host cell adhesion. Molecular microbiology, 2010. 76(3): p. 605–615.

16. Shams, F., et al., Fructose-1, 6-bisphosphate aldolase (FBA)–a conserved glycolytic enzyme with virulence functions in bacteria:‘ill met by moonlight’. 2014, Portland Press Limited.

17. Nasr, A., et al., Th-1, Th-2 Cytokines profile among Madurella mycetomatis eumycetoma patients. PLoS neglected tropical diseases, 2016. 10(7): p. e0004862.

18. Tomar, N. and R.K. De, Immunoinformatics: an integrated scenario. Immunology, 2010. 131(2): p. 153–168.

19. Hall, T., I. Biosciences, and C. Carlsbad, BioEdit: an important software for molecular biology. GERF Bull Biosci, 2011. 2(1): p. 60–61.

20. Zhang, G. and B. Fang, A uniform design-based back propagation neural network model for amino acid composition and optimal pH in G/11 xylanase. Journal of Chemical Technology & Biotechnology: International Research in Process, Environmental & Clean Technology, 2006. 81(7): p. 1185–1189.

21. Vita, R., et al., The immune epitope database (IEDB) 3.0. Nucleic acids research, 2014. 43(D1): p. D405–D412.

22. Hall, T.A. BioEdit: a user-friendly biological sequence alignment editor and analysis program for Windows 95/98/NT. in Nucleic acids symposium series. 1999. [London]: Information Retrieval Ltd., c1979-c2000.

23. Larsen, J.E.P., O. Lund, and M. Nielsen, Improved method for predicting linear B-cell epitopes. Immunome research, 2006. 2(1): p. 2.

24. Emini, E.A., et al., Induction of hepatitis A virus-neutralizing antibody by a virus-specific synthetic peptide. Journal of virology, 1985. 55(3): p. 836–839.

25. Kolaskar, A. and P.C. Tongaonkar, A semi-empirical method for prediction of antigenic determinants on protein antigens. FEBS letters, 1990. 276(1-2): p. 172–174.

26. Parker, J.M., D. Guo, and R.S. Hodges, New hydrophilicity scale derived from high-performance liquid chromatography peptide retention data: correlation of predicted surface residues with antigenicity and X-ray-derived accessible sites. Biochemistry, 1986. 25(19): p. 5425–32.

27. Andreatta, M. and M. Nielsen, Gapped sequence alignment using artificial neural networks: application to the MHC class I system. Bioinformatics, 2015. 32(4): p. 511–517.

28. Buus, S., et al., Sensitive quantitative predictions of peptide-MHC binding by a ‘Query by Committee’artificial neural network approach. Tissue antigens, 2003. 62(5): p. 378–384.

29. Nielsen, M., et al., Reliable prediction of T-cell epitopes using neural networks with novel sequence representations. Protein Science, 2003. 12(5): p. 1007–1017.

30. Peters, B. and A. Sette, Generating quantitative models describing the sequence specificity of biological processes with the stabilized matrix method. BMC bioinformatics, 2005. 6(1): p. 132.

31. Lundegaard, C., et al., NetMHC-3.0: accurate web accessible predictions of human, mouse and monkey MHC class I affinities for peptides of length 8–11. Nucleic acids research, 2008. 36(Suppl_2): p. W509–W512.

32. Abdelbagi, M., et al., Immunoinformatics prediction of peptide-based vaccine against african horse sickness virus. Immunome Research, 2017. 13(135): p. 2.

33. Patronov, A. and I. Doytchinova, T-cell epitope vaccine design by immunoinformatics. Open biology, 2013. 3(1): p. 120139.

34. Wang, P., et al., A systematic assessment of MHC class II peptide binding predictions and evaluation of a consensus approach. PLoS computational biology, 2008. 4(4): p. e1000048.

35. Wang, P., et al., Peptide binding predictions for HLA DR, DP and DQ molecules. BMC bioinformatics, 2010. 11(1): p. 568.

36. Kim, Y., et al., Immune epitope database analysis resource. Nucleic acids research, 2012. 40(W1): p. W525–W530.

37. Nielsen, M. and O. Lund, NN-align. An artificial neural network-based alignment algorithm for MHC class II peptide binding prediction. BMC bioinformatics, 2009. 10(1): p. 296.

38. Bui, H.-H., et al., Predicting population coverage of T-cell epitope-based diagnostics and vaccines. BMC bioinformatics, 2006. 7(1): p. 153.

39. Pettersen, E.F., et al., UCSF Chimera—a visualization system for exploratory research and analysis. Journal of computational chemistry, 2004. 25(13): p. 1605–1612.

40. Peng, J. and J. Xu, RaptorX: exploiting structure information for protein alignment by statistical inference. Proteins: Structure, Function, and Bioinformatics, 2011. 79(S10): p. 161–171.

41. Mohammed, A.A., et al., Epitope-Based Peptide Vaccine Against Fructose-Bisphosphate Aldolase of Madurella mycetomatis Using Immunoinformatics Approaches. Bioinformatics and biology insights, 2018. 12: p. 1177932218809703.

42. Elgenaid, S., et al., Prediction of Multiple Peptide Based Vaccine from E1, E2 and Capsid Proteins of Rubella Virus: An In-Silico Approach. Immunome Res, 2018. 14(146): p. 2.

43. Bolis, S., et al., Immunoinformatics Prediction of Epitope Based Peptide Vaccine Against Madurella mycetomatis Translationally Controlled Tumor Protein. bioRxiv, 2018: p. 441881.

44. Mohammed, A., et al., Epitope-based peptide vaccine design against Mokola rabies virus glycoprotein G utilizing in silico approaches. Immunome Res, 2017. 13(144): p. 2.

45. Albagi, S., et al., Immunoinformatics-Peptide Driven Vaccine and In silico Modeling for Duvenhage Rabies Virus Glycoprotein G. J Clin Cell Immunol, 2017. 8(517): p. 2.

46. Ahmed, O., et al., Immunoinformatic approach for epitope-based peptide vaccine against Lagos rabies virus glycoproteinG. Immunome Res, 2017. 13(137): p. 2.

47. Abu-haraz, A., et al., Multi epitope peptide vaccine prediction against Sudan Ebola virus using immuno-informatics approaches. Adv Tech Biol Med, 2017. 5(203): p. 2379-1764.1000203.

48. Sahar Suliman Mohamed, M.M.O., Samah Mahmoud Sami, Hanaa Abdalla Elgailany, Maysaa Khalid Bushara, Hadeel Abdelrahman Hassan, Ghidaa Faisal Mustafa, Amna Idris Ali, Afra Abd Elhamid Fadl Alla, Alaa Abusufian Elkabashi & Mohamed Ahmed Salih, Immunoinformatics Approach for Designing Epitope-Based Peptides Vaccine of L1 Major Capsid Protein against HPV Type 16. International Journal of Multidisciplinary and Current Research, 2016.

49. Badawi, M.M., et al., Highly conserved epitopes of Zika envelope glycoprotein may act as a novel peptide vaccine with high coverage: immunoinformatics approach. Am J Biomed Res, 2016. 4(3): p. 46–60.

50. Badawi, M.M., et al., Immunoinformatics predication and in silico modeling of epitope-based peptide vaccine against virulent Newcastle disease viruses. Am. J. Infectious Dis. Microbiol, 2016. 4(3): p. 61–71.

